# Hexadirectional modulation of EEG gamma band activity

**DOI:** 10.64898/2026.05.07.723472

**Authors:** Sein Jeung, Christopher Hilton, Christian F. Doeller, Klaus Gramann

**Affiliations:** Department of Biopsychology and Neuroergonomics,Technical University of Berlin, Germany; Max Planck Institute for Human Cognitive and Brain Sciences, Leipzig, Germany; Kavli Institute for Systems Neuroscience, Norwegian University of Science and Technology, Trondheim, Norway

## Abstract

The medial temporal lobe (MTL) houses specialised cell types supporting spatial representation in the human brain. Among these are grid cells that encode the location of the navigator, displaying a geometrically structured anchoring to the environment. While macroscopic grid-cell-like coding in humans is typically investigated using functional magnetic resonance imaging or invasive electrophysiological methods in clinical populations, we demonstrate the feasibility of using non-invasive scalp electroencephalography (EEG) to capture the characteristic six-fold modulation of high-frequency activity source-localised to the MTL. We found hexadirectional modulations of the low- and high-gamma band activity by movement direction in virtual reality. Furthermore, individual preference to use an allocentric reference frame was linked to higher hexadirectional modulation in the low gamma band and encoding a location along the putative grid axes of high gamma band predicted faster retrieval of the remembered location. The use of scalp EEG to capture grid-cell-like activity and its functional relevance sets the foundation for investigating the MTL activity in richer experimental contexts.

## Introduction

All moving animals need to represent the space around them to gain access to resources. The mechanisms of such spatial coding bear significance beyond navigation, as they reflect broader principles of how the brain structures knowledge and experience (Bellmund et al., 2018). A key component in this process is the grid cell network, housed in the entorhinal cortex (EC), that represents the structure of the external environment (Bush et al., 2015; Fiete et al., 2008; Hafting et al., 2005). First identified on the single-cell level in ambulating rodents, a grid cell is characterised by its distinct firing pattern that tessellates the space in triangular grids (Hafting et al., 2005). The geometric organisation of these firing fields leads to a hexadirectionally symmetric macroscopic signal that can be observed in human participants (Doeller et al., 2010). This property allows for translation of computational principles identified by single-unit recordings to noninvasive human neuroimaging techniques.

The characteristic hexadirectional modulation of macroscopic EC signals in human participants was originally reported using functional magnetic resonance imaging (fMRI; Doeller et al., 2010), demonstrating cyclic modulation of the BOLD signal by movement direction with six peaks in the angular space. This was followed by identification of similar effects in intracranial electroencephalography (iEEG) in the lower frequency range (4 to 8 Hz in Chen et al., 2018; 6 to 9 Hz in Chen et al., 2021; 5 to 8 Hz in Maidenbaum et al., 2018) and magnetic encephalography (MEG) in the low frequency activity (4 to 10 Hz, Convertino et al., 2023) as well as in high frequency oscillatory signal (60 to 120 Hz, Staudigl et al., 2018, with the accompanying iEEG analyses likewise showing hexadirectional modulation in the high-frequency range). However, the feasibility of applying the analysis method used in these studies to scalp electroencephalography (EEG) data has yet to be systematically evaluated. While MEG can capture information with a higher temporal resolution compared to fMRI, neither method can record brain activity during physical motion. Recent technological advances enable mobile recording of electrophysiological activity from deeper sources (Stangl et al., 2023; Topalovic et al., 2020). However, the methodology is invasive and thus limited in the accessible sample population. As a result of these methodological restrictions, evidence for recording of grid-cell-like activity in moving humans remains elusive.

Further, evidence supporting the behavioural link between the grid-cell-like activity and behaviour in human participants presents mixed results where a larger magnitude of such modulation is associated with both positive (Kunz et al., 2015; Maidenbaum et al., 2018) and negative (W. Wang & Wang, 2021) navigation performance depending on the task. Properties of grid cells indicate their role in path integration by means of tracking self-motion using the regular firing pattern as the internal spatial metric (Barry & Bush, 2012). However, the geometric organisation of grid cell activity is not relative to the self, but is known to be anchored to the external environment (Barry & Bush, 2012; Gil et al., 2018). This points to the possibility that the alignment between self-motion and the grid-like template may affect the encoding of idiothetic information to be integrated into a global representation. Testing the presence of such an effect will shed light on the functional relevance of the six-fold symmetric representation of the environment. At the same time, environment-anchored grid-cell-like representations are likely to be less salient when a non-global reference frame is in use (Peng et al., 2025). This proposition can be tested in humans by examining whether individuals who rely more on external reference frames show stronger grid-like activity.

To bridge these gaps, we utilised high-density EEG, solidifying and expanding on the findings from MEG literature (Convertino et al., 2023; Staudigl et al., 2018) that macroscopic representations in the EC can be read out in noninvasive electrophysiology recordings. We created a computerized navigation task, in which healthy participants encoded and retrieved various locations and translated between them along straight trajectories with different orientations relative to global environmental cues. The participants were grouped into egocentric and allocentric navigators based on a pretest determining their preferred use of specific spatial reference frames (Goeke et al., 2015; Gramann, 2013). Further, we examined whether the alignment between the putative grid-like template and the orientation of movement during encoding affects behaviour. Ultimately, the use of scalp EEG to this end represents a step towards the possibility of capturing human brain activity in a broader set of contexts.

Our results show that 1) hexadirectional modulation of EEG power by movement direction was identified in the low- and high-gamma band activity. Properties of the modulation aligned with previous findings on hexadirectional modulation in humans, where the putative grid orientation varied across different individuals. In addition, we identified a link between participant’s behavioural characteristics and the hexadirectional modulation. Specifically, an approach angle during learning closer to the main axes of the grid-like template derived from the high gamma band led to a faster retrieval of the remembered location. Together, these results corroborate the known findings about putative grid-cell networks in humans while adding new insights about relevant scalp EEG features and their connection to the navigation behaviour.

## Results

Forty-eight (48) right-handed participants who showed a clearly classifiable spatial reference frame proclivity in a pre-screening test (RFPT; Goeke et al., 2015) were invited to participate in the EEG experiment. They performed a spatial memory task whilst equipped with a high-density scalp EEG system. After exclusion of seven participants due to hardware issues, 41 (22 females, mean age 26.68 +-3.89 years) participants were included in the final analysis. Among the 41 participants, 19 were classified as egocentric and 22 as allocentric.

Participants navigated along consecutive straight segments presented on screen. In each trial, they were instructed to remember the location where they started and to navigate back to that location when prompted (Figure 1A). During translation of these consecutive straight segments, the portions of data with stable heading and sufficient instantaneous velocity were identified and used for further analysis (see methods for further details).

**Figure 1.**
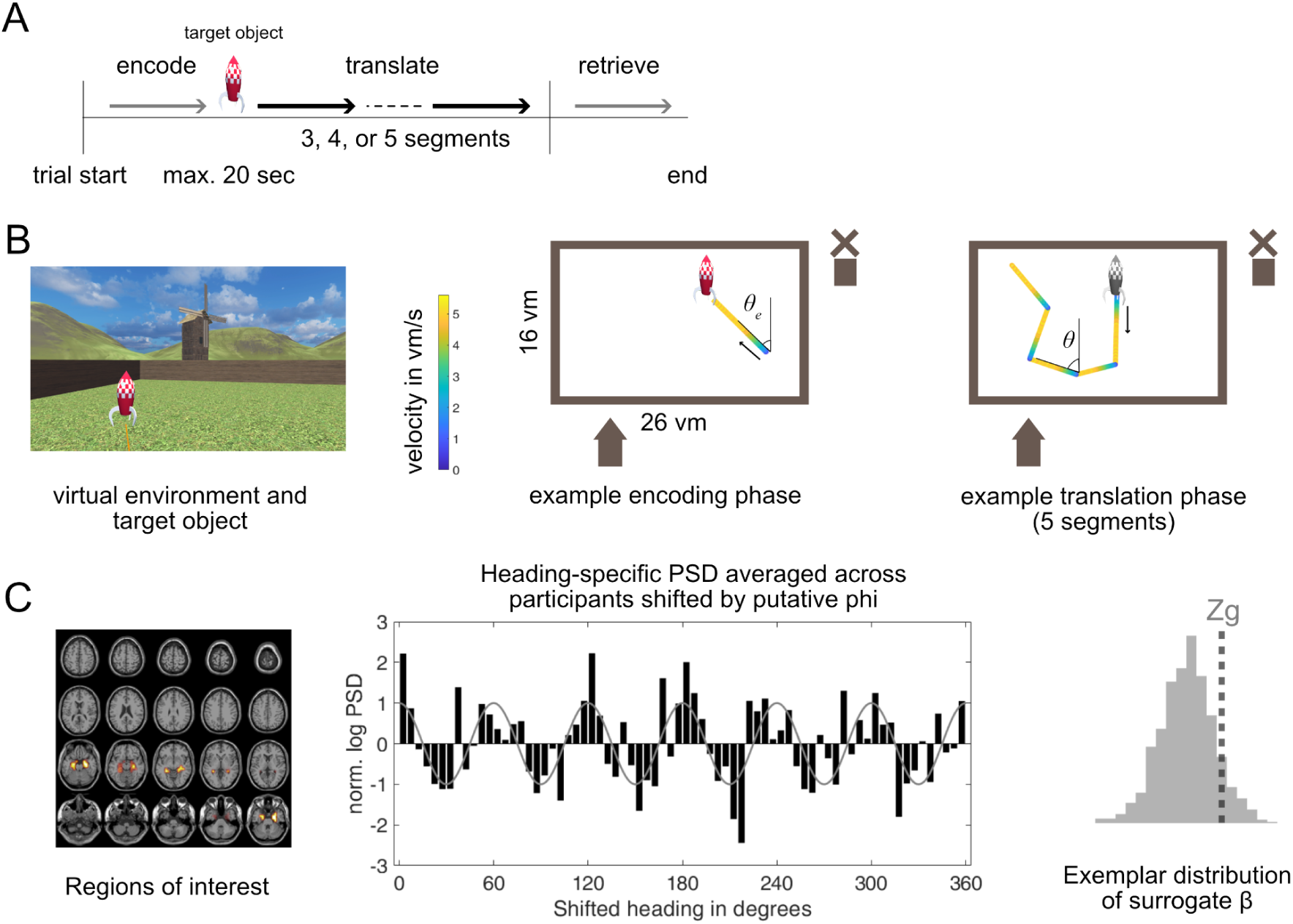
Experimental setup, navigation trajectory, and analysis logic. A. Task structure. B. Left: participant view of the virtual environment and the target during the encoding phase. Center: example participant trajectory during the encoding phase. Theta_e indicates the heading during the encoding segment, where the participant navigates to the visible target. Color coding or the trajectory indicates the movement velocity in vm/s. Right: participant trajectory during the translation phase, where the participant navigates away from the hidden target following the straight segments. Theta indicates the heading during translation of the third segment from the target. C. Left: Anatomical mask for the regions of interest-bilateral hippocampi and parahippocampal cortices. Center: heading-specific high gamma PSD in the right hippocampal voxels averaged across all participants and trials, shifted by individual putative phi. Right: exemplar distribution of surrogate betas and grid-z score (Zg).

The EEG data were preprocessed with the BeMoBIL pipeline (2022) and source-localized following the approach from Staudigl et al. (2019) with anatomical masks on bilateral hippocampi and parahippocampal cortices (Figure 1C, left). Frequencies of interest included theta (4-8 Hz), alpha (8-12 Hz), beta (12-30 Hz), low gamma (30-60 Hz), and high gamma (60-120 Hz).

### Hexadirectional modulation of human scalp-EEG signal

The hexadirectional analysis followed the method reported by Maidenbaum et al. (2019). Grid-z scores were derived from the average power in each frequency band and region of interest (ROI), to signify the magnitude of a 6-fold symmetric modulation against a surrogate distribution. Given the coarse resolution of spatial segmentation used for source localisation (AAL 3; Rolls et al., 2020), we chose the bilateral hippocampal and parahippocampal regions as broad medial temporal ROIs expected to capture the signals originating in the entorhinal cortex. The same metric was computed for 4-, 5-, 7-, and 8-fold control symmetries (Figure 2A).

**Figure 2.**
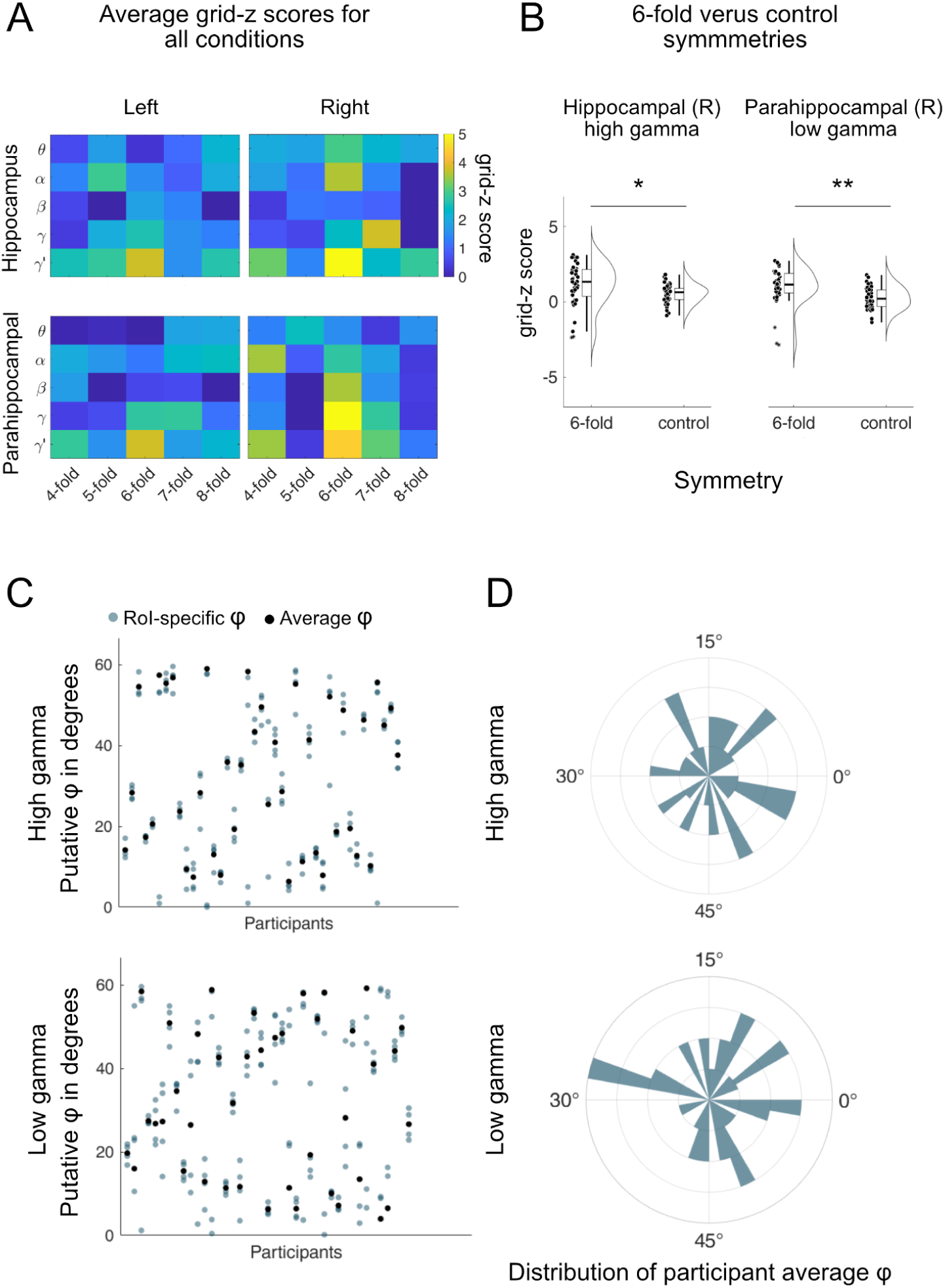
Hexadirectional modulation of high and low gamma activity. A. Summary of average grid-z values across all combinations of region of interest (ROI), symmetry (4-, 5-, 6-, 7-, 8-fold) and frequency band (θ: theta, α: alpha, β: beta, γ: low gamma, γ′: high gamma). B. Properties of putative grid orientations resulting from 6-fold symmetry analysis for low and high gamma bands. C. Participant-specific putative phis. The grey dots indicate the ROI-specific estimates and the black dots indicate the circular mean across the phis estimated for all four ROIs. D. Polar histograms of participant-specific putative phis.

In total, 20 two-tailed t-tests (4 ROIs, 5 frequency bands) were performed on grid-z scores for 6-fold symmetry. The resulting p-values were corrected using the Bonferroni method for 20 comparisons. We found six-fold symmetric modulations by instantaneous virtual heading of the participants in five combinations of frequency bands and ROIs (left hippocampus high-gamma, right hippocampus high gamma, left parahippocampal cortex high and low gamma, right parahippocampal cortex high gamma, see Table 1). For the five feature combinations for which the grid-z score was significant, we performed post-hoc comparisons of the significant symmetry against all other symmetries in the corresponding frequency band and ROI. Marginal means were estimated from a repeated measures ANOVA model with symmetry as a within-subject factor. A planned contrast (weights: −0.25, −0.25, +1, −0.25, −0.25) tested whether the cyclic modulation was not only larger than null but also larger than the average of other types of symmetries. Only the high gamma band in the right hippocampus (estimate = 0.621, SE = 0.216, t(40) = 2.877, p = 0.032) and the low gamma band in the right parahippocampal cortex (estimate = 0.776, SE = 0.207, t(40) = 3.752, p = 0.003) displayed significant differences between six-fold (Figure 2B) and control symmetries (other tests p > 0.080, see Table 1). The p-values were Bonferroni corrected for five tests.

**Table 1.** Results of t-tests and custom tests comparing 6-fold versus control symmetries.

| RoI | Band | Mean | SD | t-test |  | 6 vs. control contrast |  |  |
| --- | --- | --- | --- | --- | --- | --- | --- | --- |
|  |  |  |  | t | p | Est. | t | p |
| Hipp L | High | 0.857 | 1.451 | 3.78 | .010 | 0.314 | 1.171 | 1.000 |
| Hipp R | High | 1.153 | 1.421 | 5.20 | <.001 | 0.621 | 2.877 | 0.032* |
| Parahipp L | High | 0.934 | 1.572 | 3.80 | .010 | 0.470 | 1.952 | 0.290 |
| Parahipp R | Low | 1.027 | 1.231 | 5.34 | <.001 | 0.776 | 3.752 | 0.003** |
| Parahipp R | High | 1.035 | 1.498 | 4.42 | .001 | 0.582 | 2.365 | 0.115 |

### Properties of the putative grid orientations

We examined the properties of putative grid orientations resulting from the hexadirectional analysis of the 6-fold modulations in the high gamma band (right hippocampus) and the low gamma band (right parahippocampal cortex), based on the results of the analysis in the previous section.

Considering the relatively low spatial resolution of EEG, we tested whether the anchoring of putative grid cell activity to the environment (putative phi) extracted from one ROI predicts the putative phi estimated in other ROIs. If the hexadirectional effect from the voxels in one ROI were to propagate to other regions, the six-fold modulation from other voxels should be aligned to the same phase. Resultant vector length analysis against a surrogate distribution (Maidenbaum et al., 2018) showed that within each individual, different ROIs represented a coherent putative phi in the high gamma band (z = 2.220) but not the low gamma band (z = 1.690). Figure 2C visualizes the putative phis across different ROIs and their circular average for individual participants. We then tested whether the ROI-specific putative phis in respective frequency bands were clustered across different individuals (Figure 2D), along specific axes in the allocentric space. Neither of the two feature combinations showed clustering of putative phi within the 60 degrees space (p = 0.245 for low gamma and p = 0.438 for high gamma).

In order to estimate the spatial extent of the grid-like modulation, we binned voxel-wise PSD values according to the alignment of the headings to the putative phi. Figure 3 illustrates that the putative phi estimated for the right hippocampal ROI revealed a spatially widespread voxelwise difference between aligned and misaligned trials. In comparison, the contrast between aligned and misaligned trials was weaker for 4-fold modulation in the gamma bands and theta band 6-fold modulation. Figure 3C illustrates kernel density estimates for these voxelwise differences between aligned and misaligned headings, showing that hexadirectional modulation likely shifts the distribution of the effect across all voxels instead of showing a spatially focused effect.

**Figure 3.**
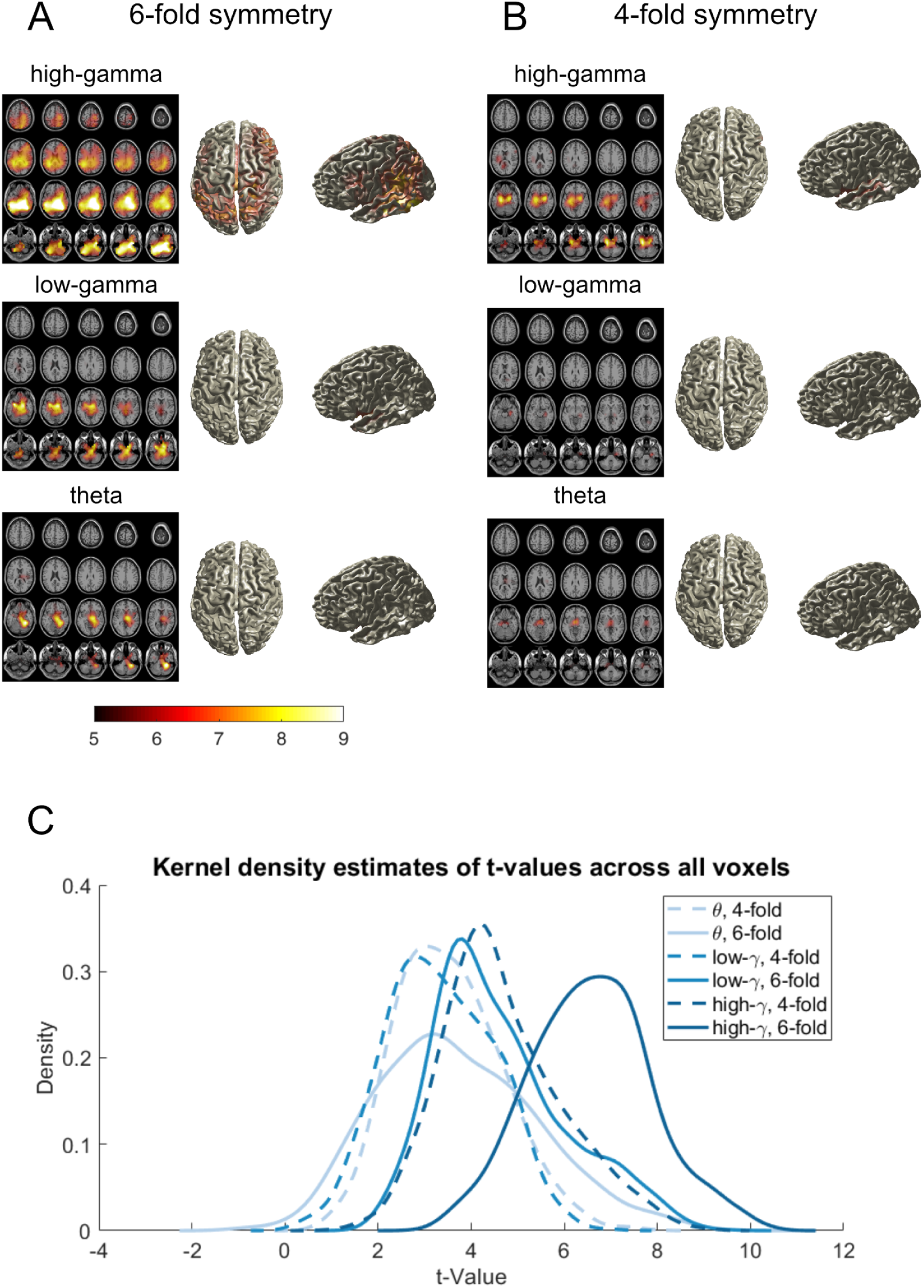
Spatial extent of hexadirectional gamma band modulation. A. Aligned voxelwise PSD - misaligned voxelwise PSD binned based on 6-fold symmetry analysis in gamma and theta bands. B. Aligned voxelwise PSD - misaligned voxelwise PSD binned based on 4-fold symmetry analysis in gamma and theta bands. C. Kernel Density Estimates for voxels in each combination of frequency band, symmetry and ROI. The spatial extent of the effect is broad as seen in the general shift of the entire distribution instead of a handful of voxels.

### Link between hexadirectional modulation and behaviour

We explored potential links between hexadirectional modulation and three behavioural variables recorded during the retrieval phase: memory scores, rotations, and trial duration. Memory score was quantified as a metric of accuracy of the retrieved location factoring in the environmental geometry (Jacobs et al., 2016), with higher scores indicating better performance. Rotation while returning to the remembered location was computed as a measure to infer a landmark-based navigation strategy (Iggena et al., 2023; Redish, 2016), with higher values indicating greater extent of horizontal rotation in the virtual environment. Duration quantified how long participants took to navigate back to the target location. The magnitude and the angular alignment with grid-like modulation were derived from the results of the ANOVA (see results above), with measures derived from high gamma confined to right hippocampal voxels and low gamma from right parahippocampal voxels.

First, we examined whether trial-by-trial measures of behaviour were predicted by the individual proclivity to use ego- or allocentric reference frames, classified using the RFPT (Figure 4A). The number of legs traversed between encoding and retrieval was included in the model as a discrete factor of three levels (3,4, and 5 legs), reflecting trial difficulty.

**Figure 4.**
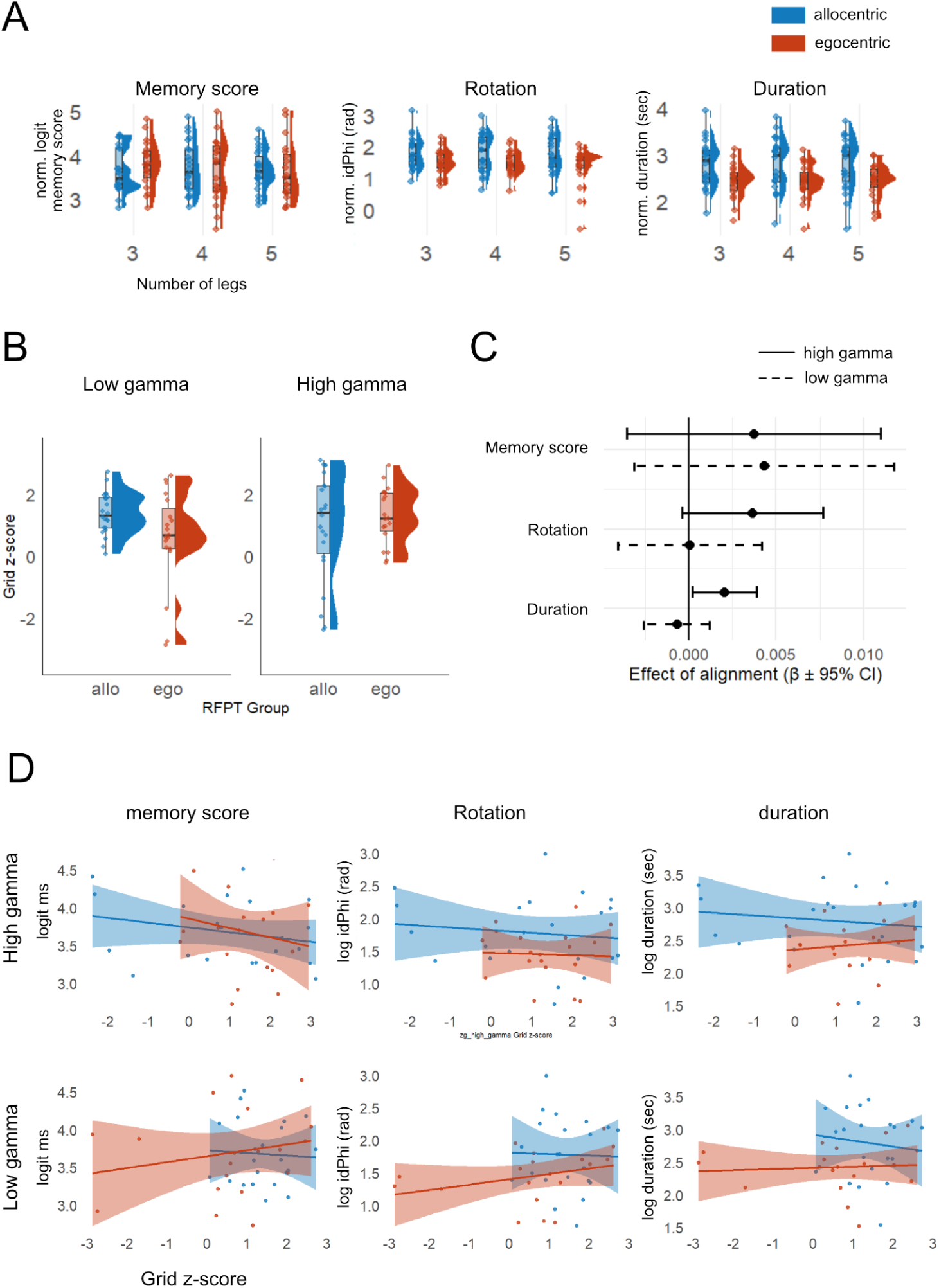
Hexadirectional modulation and navigation behaviours. Red indicates participants classified to have egocentric reference frame proclivity. Blue indicates participants with allocentric reference frame proclivity. A. Behavioural metrics of navigation performance grouped by reference frame proclivity. Participant averages of memory score, rotation, and duration grouped by the number of legs and reference frame proclivity. B. Comparison of grid-z scores between ego- and allocentric navigators, in low and high gamma bands. C. Effect of alignment between encoding heading and the individual putative phi on behaviour. Participant-specific slopes for angular distance, derived from mixed-effects models predicting behaviour, are shown on the horizontal axis. D. Relationship between grid-z scores and behavioural metrics (memory score, rotation, and duration). The vertical axes indicate the values of respective behavioural metrics and the horizontal axes indicate grid-z scores.

For memory scores, no effects of reference frame proclivity (F(1,38) = 0.01, p = .940), number of legs (*F*(1,2906.3) = 0.43, *p* = .648), or their interaction were found (*F*(1,2906.3) = 1.08, *p* = .341, allocentric M = 3.68, SD = 0.51, egocentric M = 3.70, SD = 0.64; mean values have been logit transformed; see Methods for derivation of memory scores).

We observed less rotation (measured as idPhi; see Methods) in egocentric navigators (M = 1.45, SD = 0.48) compared with allocentric navigators (M = 1.78, SD = 0.58) revealing a significant difference in landmark-based navigation behaviour (*F*(1,38) = 4.39, *p* = .042). Moreover, the number of legs had a statistically significant effect on rotation (F(2,2905.1) = 4.13, p = .016). Post-hoc pairwise comparisons with Bonferroni adjustment indicated that participants showed more rotations in trials with 5 legs compared to 3 legs (estimate = 0.11, p = .033) and compared to 4 legs (estimate = 0.11, p = .046) but there was no difference between 3 and 4 legs (estimate < 0.01, p = 1.00). No interaction between preferred reference frame and number of legs was identified (*F*(1,2905.1) = 0.27, p = .763).

For *duration*, a significant effect of reference frame proclivity emerged (*F*(1,38) = 6.14, *p* = .018), with egocentric participants completing trials faster (M = 2.43 seconds, SD = 0.37) than allocentric participants (M = 2.80, SD = 0.55). No effects of number of legs (*F*(1,2906) = 0.06, p = .946), nor an interaction between reference frame proclivity and number of legs were observed (*F*(1,2906) = 0.17, p = .844).

Next, we tested whether reference frame proclivity was related to the magnitude of hexadirectional modulation. We found a significant effect of reference frame proclivity such that egocentric participants had lower low-gamma grid scores compared with allocentric participants (F(1,39) = 4.69, p = .037) . However visual examination of Figure 4B indicated that the effect may be driven by two extreme values. Consequently, we performed IQR based outlier detection which indeed removed these data points and the effect was no longer significant (*p* = .166). For high gamma, no effect of reference frame proclivity on grid z-scores was identified (F(1,39) = 0.48, *p* = .491).

We then tested the effect of movement direction relative to the phase of hexadirectional modulation on navigation behaviour. This relationship between movement direction and the phase of hexadirectional modulation was quantified as the angular distance between the orientation of the encoding segment and the nearest of the six orientations for which the PSD estimates peaked (corresponding to the putative phi). The angular distance derived from high gamma band activity predicted the duration of the retrieval phase (F(1,2908) = 4.74, *p* = .030, β = 0.0020 ± 0.0009 SE, Cohen’s *d* = 0.08; Figure 4C, right panel). The positive relationship indicates that the higher the angular distance between the putative phi and encoding orientation was, the longer it took for the participants to retrieve the remembered location. The alignment of low or high gamma putative phi and the encoding orientation was not significantly related to any of the other variables (all ps > .076).

Building on the behavioural analyses above, we next examined whether EEG-derived gamma power predicted navigation behaviour. Neither low- nor high-gamma power predicted any of the behavioural variables (all *p* > .125, Figure 4D).

## Discussion

Our results confirm observations from the iEEG, MEG and fMRI literature showing that humans display grid-cell-like activity in medial temporal areas. The hexadirectional modulation observed in the low (60-90 Hz) and high gamma (90-120 Hz) bands is consistent with MEG findings (60-120 Hz; Staudigl et al., 2018). Furthermore, the estimated phases of putative grid cell systems did not cluster across participants, despite the asymmetry of the virtual environment. Notably, the phase of hexadirectional modulation of the gamma band activity was associated with the duration of retrieval of a remembered location.

The pattern of lateralisation and frequency-band effects offers methodological insight into how grid-cell-like coding can be captured using scalp EEG. The right-lateralized modulation observed in our data suggests hemispheric specialisation in grid-cell-like processing, reflecting the predominant role of the right medial temporal lobe in spatial navigation (Miller et al., 2018). Localisation of hexadirectional modulation to the left MTL has been reported when the task has no navigation component (Staudigl et al., 2018), while right lateralisation is more reliably observed in navigation contexts (Convertino et al., 2023; Doeller et al., 2010; He & Brown, 2019). In addition, we did not observe a hexadirectional symmetry effect in the lower frequency ranges, unlike previous electrophysiology studies reporting hexadirectionally symmetric modulation of theta band activity (Chen et al., 2018, 2021; Convertino et al., 2023; Maidenbaum et al., 2018). Electrophysiologically measured scalp theta activity appears to be best characterized when time-locked to movement onset (Convertino et al., 2023), whereas we analyzed time windows after the initial acceleration phase of the movements. The known links between high-frequency neural activities and theta oscillations (Canolty et al., 2006; N. Wang et al., 2025; White et al., 2012) suggest that the observed modulation of gamma band activity serves as a practical electrophysiological marker potentially manifesting theta-gamma coupling.

The distribution of the orientations of the putative grid-like template estimated for different individuals did not statistically deviate from a uniform distribution (Figure 2D), showing that the grid was not anchored to the geometric boundary of the environment. Previous research provides mixed evidence in this regard, where some findings show boundary anchoring of grid orientation (He & Brown, 2019; Krupic et al., 2015), whereas the distribution of putative grid-like templates is shown to be uniform in the absence of a strongly polarizing cue (Julian et al., 2018; Maidenbaum et al., 2018). The rectangular environment in our task elongated the navigable space along a single axis, but the presence of multiple distal cues, including three landmark objects and a visually rich skybox, may have led to between-individual variance in orientation anchors.

Individual variance in orientation anchoring may be further explained by preferences in reference frame use for spatial encoding. On the group level, individuals with an egocentric reference frame proclivity showed a lower magnitude of hexadirectional modulation than allocentric navigators, suggesting systematic differences in the expression of grid-like coding as a function of preferred spatial reference frame. This pattern is in line with the notion that an allocentric reference frame supports the use of stable distal cues for spatial representations. This finding should be treated cautiously, as the effect did not survive removal of data points with extreme values. Importantly, converging behavioural evidence supports a related differentiation between reference frames. Egocentric navigators displayed less rotation behaviour, indicating a reduced reliance on landmarks (Iggena et al., 2024). In addition, the faster retrieval times in the egocentric group are consistent with accounts suggesting that egocentric, route-based navigation can support more direct action selection, whereas allocentric navigation rely on cognitive map-based processing and may involve additional computational steps such as viewpoint transformations and prospective simulation (Hartley et al., 2018; Schmidt et al., 2013). The observed grid-z scores were in the negative range in the egocentric participants with extreme values, alluding to a systematic deviation from the surrogate distribution in the direction opposite to the prediction. In our hexadirectional analysis it is assumed that the grid-cell-like representation is stably anchored to an external reference frame across trials. Our approach would result in a lower magnitude of modulation in egocentric navigators if they utilise a different spatial anchor, such as specific task-relevant cues defining the reference frame (Clark & Nolan, 2024; Peng et al., 2025), and even a negative index for modulation score if there is a systematic switching in the choice of anchor between different portions of the experiment. Such dynamic mapping between the grid cell system and the environment would be driven by individual experience and behaviours (Ginosar et al., 2023) which is supported by the lack of a global alignment of grid orientations across participants in our data (Figure 2D).

We observed a link between the magnitude and phase of gamma-band modulation and navigation behaviour, which suggests that hexadirectional modulation may serve a functional role in the navigation process, rather than representing a mere byproduct of neural processing. Specifically, the duration of the retrieval phase was predicted by the relationship between heading direction during encoding and the putative grid orientation. The extremely small effect size (Cohen’s *d* = 0.08) highlights that while the influence of approach direction on retrieval is detectable, it is subtle. Nonetheless, on the theoretical level, it supports the idea that grid cells incorporate proximal, action dependent perception of the space (Ginosar et al., 2023). Taken together, our findings align with the view that grid cells may anchor to multiple local reference frames, not only to a single global reference frame (Peng et al., 2025), and demonstrate the possibility to link sub-cortical neural activity to individual behavioural dynamics.

While these results provide novel insight into the functional relevance of grid-like representations during navigation, several methodological factors should be considered when interpreting the spatial distribution of the observed effects. We observed a rather broad spread of hexadirectional modulation in the low and high gamma bands, as visualised in Figure 3. We used a standard template head model rather than individualised anatomical scans for source localisation, which is known to negatively affect the localisation accuracy of scalp EEG data (Klamer et al., 2015). Whether the observed effect reflects the physiological characteristics of entorhinal activity that propagate to other regions (e.g., prefrontal cortex, Convertino et al., 2023; Park et al., 2021) or is simply an artefact of limited spatial precision of the source localisation method remains unclear.

As different neuroimaging methods have been found to reveal distinct aspects of the same cognitive process (Robitaille et al., 2010), fortifying this junction through simultaneous fMRI-EEG or iEEG-scalp EEG is expected to open new opportunities to better characterize grid-cell-like dynamics in humans. At the same time, our findings lay the groundwork for future applications using mobile EEG in naturalistic and ecologically valid setups. This would better approximate the behavioural context of prior animal studies, as mobile EEG enables the investigation of interdependent neural and behavioural dynamics during motion (Debener et al., 2012; Gramann et al., 2011; Jungnickel et al., 2019; Makeig et al., 2009).

## Methods

The scripts and documentation for trajectory creation, preprocessing and analysis are publicly available on Github (https://github.com/sjeung/Hexa-trajectory for trajectory creation scripts, https://github.com/sjeung/Hexa-Analysis for preprocessing and analysis). The raw data set is publicly available on OpenNeuro (doi:).

### Participants

41 right-handed participants were included in the analysis, consisting of 19 egocentric navigators (F = 10, age 22 to 35 years old, M = 26.95, SD = 3.50) and 22 allocentric navigators (F = 12, age 20 to 36 years old, M = 26.45, SD = 4.26). Originally, 48 participants with no known history of neurological condition were recruited through the participant portal of the Institute of Psychology and Ergonomics at the Technical University of Berlin. Seven participants were excluded: six due to EEG hardware issues, and one due to excessive simulator sickness. The protocol was reviewed and approved beforehand by the ethics committee of the institute (application number GR_13_20191008). All participants provided written informed consent and received monetary reimbursement of 10 euros per hour or one course credit per hour. Participants additionally read and signed the written forms about the hygiene protocol implemented due to the outbreak of COVID-19. The duration of the experiment in the lab was between 3 and 3.5 hours including the formal instructions and preparation time for EEG recording.

Prior to the lab appointment, an online pretest was administered to categorise the participants according to their preferred use of reference frames using the tunnel paradigm (Goeke et al., 2015; Gramann et al., 2005). We used a version of the task hosted on Labvanced (Link: https://www.labvanced.com/player.html?id=1504), an online platform for computerised experiments (Finger et al,. 2017). Among 24 trials, at least 75% of ego or allocentric responses categorized them as either. Individuals with unclear preference or switching reference frames were not invited to the main experiment. Participant groups were stratified according to sex (M and F) and reference frame proclivity (egocentric and allocentric), with 12 participants in each of the four subgroups. Fourteen of the online participants with unclear reference frame proclivity or whose strategy was categorised to belong to a fully recruited subgroup were not invited to the lab session and received partial reimbursement for participation in the pretest.

### Hardware setup

Participants were equipped with a wireless BrainProducts MOVE system. 128 active gel-based wet electrodes (BrainProducts actiCAP slim) were mounted on a BrainProducts actiCAP snap cap with a custom equidistant layout approximating 10-5 system locations (Oostenveld & Praamsta, 2001). The locations of sensors relative to the anatomical landmarks were digitised using a 3D position measurement system (Polaris Vicra, NDI, Waterloo, ON, Canada). Two electrodes were placed under the eyes to capture activity originating from vertical eye movements. The EEG stream was continuously recorded throughout the sessions, with a nominal sampling rate of 1000 Hz. Data was referenced to an electrode located adjacent to the standard 10-20 position F3 and a ground electrode located near F4. BrainAmp DC amplifier was used with 0.016Hz high-pass (first-order with 6 dB/octave) and 500 Hz low-pass (fifth-order Butterworth filter with 30 dB/octave) online filters. High-viscosity electrolyte gel (SuperVisc) was used to establish contact between the scalp and the electrodes, with the target Impedance level below 10 KOhm.

Participants performed the task standing in front of a wall-mounted 43-inch monitor with a refresh rate of 60 Hz. The height of the monitor was adjusted to the individual preferences. The virtual environment was created using Unity 3D (version 2018.2.8f1). Participants controlled the virtual camera using the arrow keys on a keyboard. Translation speed was set to 1.4 vm/s (virtual metres per second). Maximum translation speed was set to 3.5 vm/s. Acceleration was linear with the rate of 0.07 virtual metres per frame (1/60 seconds). Rotation speed was fixed at 60 degrees per second. The virtual motion stream consisting of position in Cartesian coordinates and orientations in quaternions were sampled at 60 Hz.

Both motion and EEG data were streamed via the LabStreamingLayer (LSL, Kothe et al., 2024) and written in extensible data format (XDF) together with sample-by-sample time stamps. Streaming of the motion data was achieved using the LSL4Unity plug-in (https://github.com/labstreaminglayer/LSL4Unity) where the data was transmitted via WiFi from the experiment PC (Zotac) to the recording PC. EEG was transmitted to the amplifiers using the BrainProducts MOVE system and streamed to LSL via the BrainVision RDA interface on the recording PC.

### Experimental task

The experiment started with the passive viewing task, in order to induce a comparable initial exposure to the virtual environment across the participants. The camera initially showed a static view for 30 seconds and then translated along all four edges of the rectangular arena, revealing the major visual features such as the boundaries and landmarks (see SI Figure S1 for images). The participant was then instructed to use the arrow keys on the keyboard to translate (up-down for forward-backward translation) and rotate (left-right) in the subsequent baseline walking task as well as in the main task. The baseline walking task consisted of passive following of waypoints that appeared sequentially in 36 different locations within the arena. An orange coloured straight line was presented on the ground to guide the participant to walk straight to the subsequent waypoint. The baseline task was followed by a practice session, in which the participant performed an example block (9 trials) while being given oral instructions from the experimenter explaining the trial structure.

A trial started with the participant learning the location of a new target object that appeared at a predefined location within the arena (see Figure 1B). They were guided by an orange line on the ground to walk straight to the target. During this encoding phase, participants were asked to first adjust their virtual heading before translating and maintain the heading while translating. Once they reached the target, they were given up to 20 seconds to look around and memorise the location. They were able to terminate this encoding phase earlier by pressing the spacebar. The following translation phase consisted of 3, 4, or 5 straight segments. The number of the segments in the navigation trajectory were balanced and randomised within each block. Upon reaching the end of the last straight segment, they were instructed to navigate back to where they remembered the target object was, and then press the spacebar (retrieval phase). The real target model appeared and participants were instructed to navigate back to the true location of the target object. When they reached the target object, the next trial started, repeating the same structure with the target object appearing elsewhere in the arena.

In order to evenly sample movement directions in the 360 degrees angular space over the course of all navigation phases in the experiment, an algorithm was constructed that sequentially samples directions from a pool and randomises the lengths (see SI for details). First, a pool of orientations was created, containing 72 unique orientations between 0 and 360 degrees in steps of five degrees. Two restrictions were imposed in each sampling step in order to create a seamless navigation trajectory within the geometrically bounded environment. 1) The turn angle had to be smaller than 120 degrees, and 2) the next sampled point had to be within the arena. For creating the translation segments, a similar procedure was applied to the pool of 72 orientations. For each trial we selected three, four, or five orientations (for the target location of that trial), and the length of each translation segment was sampled uniformly from five to seven virtual meters. In addition, the above restrictions on the turn angle and bounded area applied. This way we ensured that the angular space is evenly sampled within the spatial constraints of the rectangular area.

The experiment consisted of 8 blocks, each including 9 trials. Between every pair of blocks the participant had an option to take a brief, self-paced break. The baseline task was presented three times, once at the beginning, the second time between the fourth and the fifth blocks, and lastly after the final block.

### Analysis

Matlab 2021a (The Mathworks Inc.) was used for preprocessing and hexadirectional analysis of the EEG data. Detailed description of preprocessing parameters can be found in using the ARTEM-IS report (Styles et al., 2021) in the SI and the Github repository containing analysis scripts (https://github.com/sjeung/Hexa-Analysis).

### EEG preprocessing

The data set was first structured according to the Brain Imaging Data Structure (BIDS, Gorgolewski et al., 2016) following the extensions for EEG (Pernet et al., 2019). The BIDS-formatted EEG data were then imported into EEGLAB (Delorme & Makeig, 2004) as SET files, using the BeMoBIL pipeline (Klug et al., 2022). Here, both the motion and EEG data streams were resampled to 500 Hz and synchronised by matching the time stamps provided by LSL (Kothe et al., 2024).

Next, the non-experimental portions of the continuous recording including breaks and practice trials were removed and the resulting data segments were concatenated back together as a continuous data structure. The BeMoBIL pipeline (Klug et al., 2022) was used for further preprocessing steps. First, bad channels were detected using the *clean raw data* EEGLAB plug-in, using the outlier detection algorithm in the PREP pipeline (Bigdely-Shamlo et al., 2015). On average, 10.171 ± 1.327 (mean ± SEM) channels per participant were detected as bad. The bad channels were removed and reconstructed by means of spherical interpolation. All channels were then average-referenced. Electrical noise from power lines was removed at a predefined frequency of 50 Hz using ZapLine (de Cheveigné, 2020).

In order to separate the channel-level signal into independent components, the Adaptive Mixture Independent Component Analysis (AMICA, Palmer et al., 2011) was used. For optimal decomposition of the signal, two processing steps were applied to AMICA input data. The data was filtered with the high-pass cutoff of 1.75 Hz. Automatic rejection of data points was performed using the built-in algorithm within the AMICA software, with the number of rejections parameter set to 10 and the sigma threshold to 3 (8.880 ± 0.417 % of samples rejected per participant). The number of models was set to one, with a maximal number of iterations of 1000.

The spatial filters for independent components were copied back to the data before high-pass filtering and automatic sample rejection steps. Then, another high-pass filter with a cutoff of 0.1 Hz was applied. The independent components were classified using ICLabel (Pion-Tonachini et al., 2019). The components exceeding the threshold of 80% in eye, heart, line, channel, or muscle category were projected out (15.122 ± 0.884 components were rejected per participant).

### Motion preprocessing and data segmentation

After BIDS conversion following the extension for motion data (Jeung et al., 2024), the motion data was resampled to match the sampling rate of the resampled EEG data (500 Hz). The time synchronisation between EEG and motion data was achieved using the time stamps created by LSL. Motion data was then formatted into .set files to facilitate temporally synchronous joint analysis together with EEG. The data were filtered with a low-pass cutoff of 6Hz. Quaternion values representing orientations were converted to Euler angles to construct the yaw time series quantifying rotations around the vertical axis in the virtual environment (e.g. looking left and right).

We extracted data segments with sustained heading for more than 500 milliseconds. During the navigating portion of a trial, a moving window was constructed so that segments were extracted based on velocity and angular deviation threshold. Data segments with sustained speed of over 2 virtual metres per second and maximal absolute angular deviation from the angular average of the orientation during the time window below 5 degrees were selected and included in the analysis. On average, 532.44 epochs were generated per participant (SEM = 8.48). For source localisation, an extended time window of 2.5 seconds was used to ensure that low frequency activities were robustly estimated. For hexadirectional analysis, only the 500 ms with 200 ms buffer before and after this time window was used.

### Source localisation

Source localisation was performed using FieldTrip toolbox (Oostenveld et al., 2011) following the approach described in an MEG study (Staudigl et al., 2018) and expanding it on the frequency domain. A template MR image included in the FieldTrip toolbox was used to warp the individually measured electrode locations onto the template image. Four ROIs were created including the right and left hippocampi and parahippocampal cortices. To construct the left and right medial temporal ROIs, we used anatomical masks including the labels ‘Hippocampus_L’, ‘ParaHippocampal_L’ and ‘Hippocampus_R’, ‘ParaHippocampal_R’, respectively, based on the Automatic Anatomical Labelling atlas in Montreal Neurological Institute space (AAL 3; Rolls et al., 2020).

The procedure for extracting voxel-wise power was largely based on Staudigl et al. (2018). However, no participant-specific MR scans were available in this study and no parcellation of the entorhinal mask was implemented. The cross-spectral density (CSD) was derived in order to construct the spatial filters. The segments were fourier-transformed within the frequency bands of interest with 2 Hz spectral smoothing using a multitaper approach. Frequency bands were defined as follows: theta (4–8 Hz), alpha (8–14 Hz), beta (14–30 Hz), low gamma (30–60 Hz), and high gamma (60–120 Hz). The frequencies of interest (FOIs) were logarithmically spaced between 4 and 128 Hz in steps of 0.125 in log-2 scale and then binned into the five frequency bands. The cross-spectrum was regularised prior to matrix inversion by loading the diagonal of the matrix with 5% of the average sensor power. For DICS beamforming (Gross et al., 2001), the center frequency of each band was calculated as the arithmetic mean of its lower and upper edges. The closest matching FOI from the precomputed frequency list was selected for source reconstruction (5.96, 10.89, 21.89, 45.25, 90.51 Hz for theta, alpha, beta, low gamma, and high gamma bands, respectively). The dipole orientations were fixed, such that only the largest of the three dipole directions per spatial filter was kept.

Then, the sensor level data was projected into source space by multiplying it with the resulting spatial filter, allowing for further analysis to be conducted in virtual sensor space. Time-domain activity of the virtual sensors were decomposed again into frequency domain to span the entire range of each frequency band of interest using a sliding window Fourier transform. Spectral power was estimated using a fixed time window of 250 ms, applied in steps of 50 ms from –500 to 1000 ms relative to the event of interest. A Hanning taper was applied to each time window to minimize spectral leakage. The FOIs were identical to those used for CSD computation and were analyzed individually, each using a 250 ms window length. The resulting power values were averaged across voxels within the ROI, across FOIs within the band, and then across the time points between 0 and 500 ms. The segment-wise power estimates were retained for further analysis for each ROI and frequency band combination.

### Hexadirectional symmetry analysis

To examine whether the PSD during translation segments were six-fold symmetrically modulated by the heading, a hexadirectional GLM method originally described in Doeller at al. (2010) was adopted by integrating the steps described in Maidenbaum et al., 2018 and Staudigl et al., 2018.

For each participant and for each combination of ROI and frequency band, vectors of the instantaneous headings and the average PSD during translation segments were computed following the procedures described above. The PSD values were first log-transformed and the segments with log-PSD values exceeding 1.5 × the interquartile range (IQR) of the distribution above the third quartile or below the first quartile were excluded from hexadirectional symmetry analysis. In the high gamma band in the right hippocampal voxels, 4.40 % of segments were removed, resulting in 509 +-8.17 segments per participant.

Translation segments were randomly divided into five sets of roughly equal size for five-fold cross-validation. For each fold, one set was held out as the test set, and a GLM predicting the log-transformed PSD values was fitted to the remaining segments. To embed the hexadirectional symmetry, the heading values (𝜃) were multiplied by six. Regressors for cosine and sine (𝜷1 and 𝜷2) of 6*𝜃 were fitted to predict the PSD (Eq. 1). Heading alignment angle (φ) was computed by taking the inverse tangent of the ratio of 𝜷2 to 𝜷1 and dividing by six (eq. 2). The procedure was repeated across all five sets, producing a set of five putative φ values for the five iterations of cross-validation.

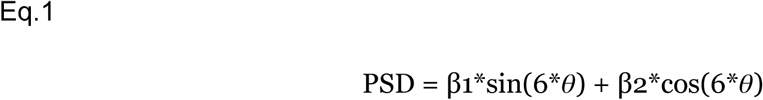

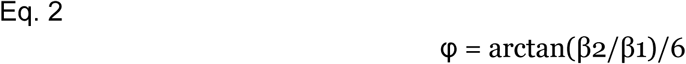

Next, we fitted another GLM (GLM II) using as regressor the cosine of heading angles after shifting to align to the estimated putative φ and scaling six-fold, i.e. cos(symmetry*(𝜃-φ)) following the procedure described in Maidenbaum et al. (2019, see eq. 3). A positive regression weight would indicate mouldation of neural activity by the six-fold rotational symmetry (Figure 1C middle panel demonstrates aggregation of all trials using the same shifting procedure, although the main analysis was implemented separately for different individuals). This operation was repeated five times for all segments, to reflect different φ estimates from 5-fold cross validation. The five 𝜷i values resulting from this GLM were averaged to yield 𝜷-bar.

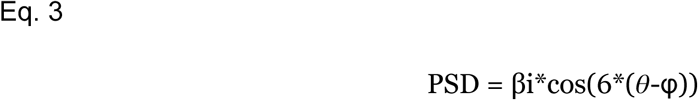

In order to control for spatially confounding properties of PSD values inflating the beta values, we fitted the cos(symmetry*(𝜃-φ)) above to shuffled data sets created by randomly assigning PSDs to headings. Like in the GLM II step, this operation was performed on five different sets of data segments. The resulting betas were averaged across the five sets. This was repeated 1000 times, to create a surrogate distribution of such 𝜷 values. We computed the z score of the 𝜷-bar values against this distribution, yielding the **grid-z** score, Zg, an estimate of the magnitude of cyclic modulation. For the combinations that showed a significant geometric symmetry, we used a custom contrast (main fold vs average of 4-, 5-, 7-, and 8-folds) to compare the symmetry of interest against all other types of symmetry.

### Properties of putative phis

We computed resultant vector length to confirm that within a single participant, the headings estimated in different ROIs were significantly similar. The surrogate distribution for resultant vector length was created following the procedure described in Maidenbaum et al., 2018. We performed a rayleigh test (using the circstat toolbox; Berens, 2009) in order to examine whether the putative grid orientations are clustered across participants.

### Whole-brain plots

In order to visualize the spatial spread of the grid-cell-like activity found in the right hippocampal high gamma band and the right parahippocampal low gamma band, we first categorized the epochs into aligned bins (absolute difference between segment theta and putative phi smaller than 15 degrees) and misaligned bins (absolute angular difference between theta and putative phi larger than 15 degrees). The virtual channel PSDs were estimated for each voxel. In a manner consistent with the main hexadirectional symmetry analysis, the PSD values were first log-transformed and IQR removal was performed on the trial level, discarding trials with the average log PSD across all voxels above the IQR thresholds estimated for each participant. In order to factor in the scale difference between frequency bands, normalisation was performed using the mean and standard deviation computed from all voxels, resulting in z-scored PSDs of aligned voxels and misaligned voxels. The voxelwise difference between z-scored aligned and misaligned PSDs were saved for each participant. Then, a right-tailed t-test was performed on these voxelwise differences in order to quantify how much higher the power was in aligned epochs compared to misaligned ones in each voxel. The resulting t-values were visualized on template MR scans (Figure 3A and 3B). Kernel density of the voxelwise t-values was estimated in order to visualise the spatial specificity of this effect across voxels (Figure 3C).

### Link to behaviour

The link between individual grid-z score and the behavioural score was examined. Memory score was used as a metric of performance that takes the bounded geometry of the environment into consideration (Bellmund et al., 2020; Jacobs et al., 2016). In order to compute the memory score of a trial, we first spawned 1000 random points within the rectangular area. For each iteration, random x and y coordinates were sampled from a uniform distribution along the range of respective axes. The Euclidean distance of this random point and the true location of the target was compared with the Euclidean distance between participant response and the target location. The memory score was defined as the proportion of random points that had a larger Euclidean distance from the target than the participant response, ranging from 0 to 1 (with 1 representing perfect performance), with the chance level at 0.5.

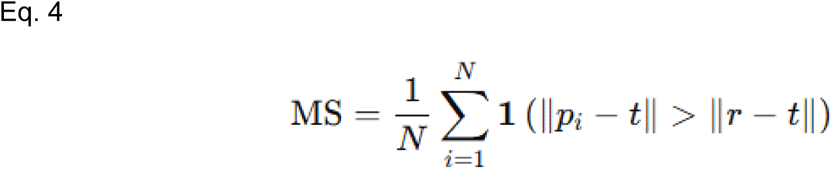

Where r is the response, t is the target, and Pi (between 1 and N) is the set of N = 1000 uniformly sampled random points and 1(.) the indicator function equal to 1 if the condition is fulfilled.

Next, we computed rotation as integrated absolute angular velocity (idPhi; Papale et al., 2016; Redish, 2016; Schmidt et al., 2013).

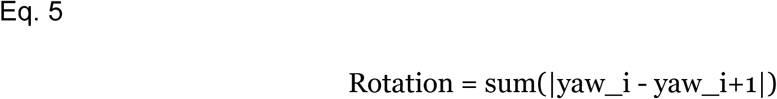

The duration of the retrieval phase was defined as the time window between the onset of the prompt to navigate back to the remembered target location and the participant response registered by a key press.

The magnitude and phase of the grid-like activity for individual participants were computed separately for the two ROI and frequency band combinations that showed heightened hexadirectional symmetry compared to others, on the basis of the results of the main hexadirectional symmetry analysis (t-tests on z-score and post-hoc ANOVA). High gamma grid-like activity was thus characterized in right hippocampal voxels and low gamma in right parahippocampal cortex voxels. The angular alignment during encoding was characterized for each trial by computing the distance of the heading during encoding (𝜃e, see Figure 1B middle panel) from the nearest grid-like template axis, ranging between 30 and 0 degrees (see Eq. 6).

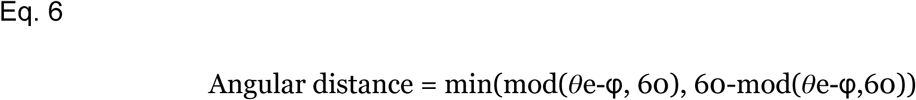

Behavioural analysis was performed in R (R Core Team, 2024, version 4.4.2) using the lme4 package for mixed effect models (LME; Bates et al., 2015). P-values were obtained using Satterthwaite’s method to estimate degrees of freedom. Model effects were assessed using the ‘anova’ function on the modelled data, and post-hoc pairwise contrasts for significant multi-level factors were performed using the emmeans package (Lenth et al., 2024). We first tested whether reference frame proclivity affected the magnitude of grid-like modulation on EEG activity.

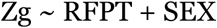

We used LME models to test whether our grouping variables or experimental manipulation were predictive of any behavioural measures.

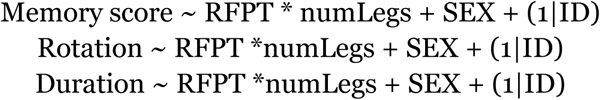

We examined the participant-specific Zg to test whether the magnitude of the hexadirectional modulation affected navigation behaviour.

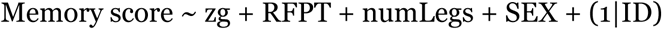

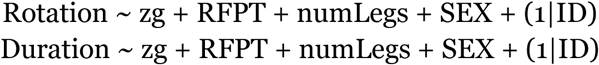

We examined the alignment with the individual putative phi during encoding as a continuous predictor variable, in order to test whether trials whose headings were aligned to the putative phi would be linked to navigation behaviour.

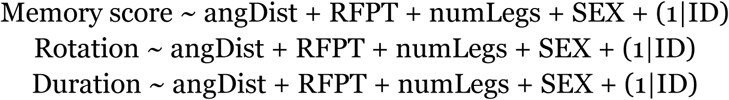

## Acknowledgements

This project received initial support from the International Office of Technische Universität Berlin and additional funding from the Department of Biopsychology and Neuroergonomics. We acknowledge Benjamin Paulisch, Yiru Chen, and Timo Berg who supported data collection.

## Declaration of interests

The authors declare no competing interests.

## Declaration of generative AI and AI-assisted technologies in the writing process

During the preparation of this work the authors used ChatGPT (OpenAI) for language editing and drafting assistance. After using this service, the authors reviewed and edited the content as needed and take full responsibility for the content of the published article.

## Supplementary Materials

### Virtual environment

All external assets for the Unity 3D project were obtained from the Unity Asset Store under the Standard Unity Asset Store EULA. The sky box was part of AllSkyFree Set (Epic BlueSunset by rpgwhitelock). The windmill model was part of Medieval Village Environment pack (by Asset Maiden) and the house model was shared as OLD HOUSE (by GBAndewGB). The target rocket model was part of ToysPack by FunFant.

**Figure S1.**
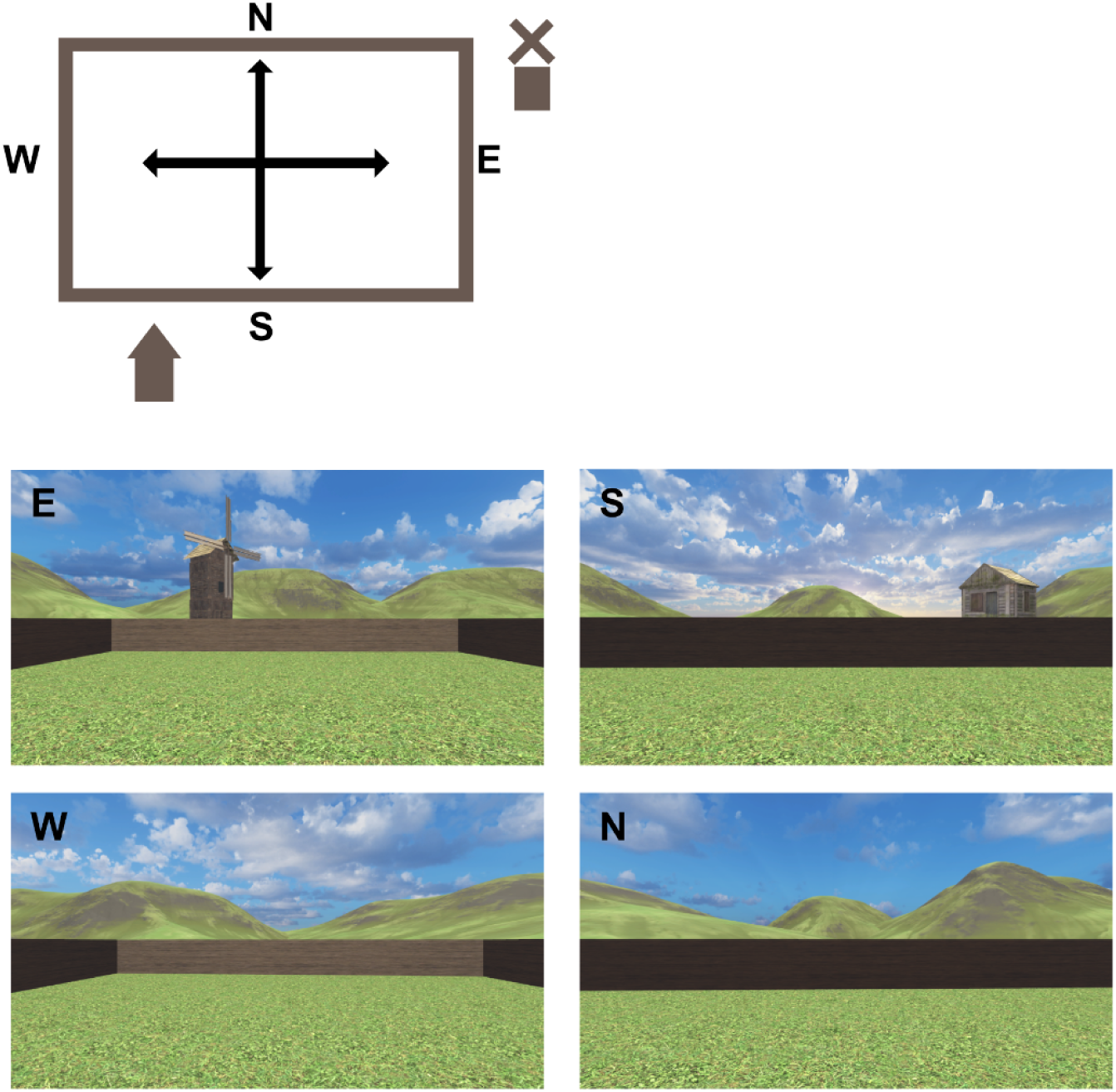
The virtual environment from four different views. The initial viewing task presented the scenes translating along the walls in the order of E-S-W-N.

### Spatial characteristics of presented trajectories

First, a pool of orientations was created, containing 72 unique orientations between 0 and 360 degrees in steps of five degrees. Starting from the center of the environment (coordinates 0,0), a randomly sampled sequence of 72 orientations was generated, with the length of each segment sampled from a uniform distribution within the interval between five and seven virtual meters. These served as the 72 encoding segments, connecting all target locations, as the participant always returned to the target location at the end of a trial before encoding the next one. Two restrictions were imposed in each sampling step in order to create a seamless navigation trajectory within the geometrically bounded environment. 1) The turn angle had to be smaller than 120 degrees, and 2) the next sampled point had to be within the arena. The orientations violating these restrictions were assigned the sampling probability of zero. The algorithm was recursively constructed such that it started over whenever it hit a dead end with no more possible moves.

For creating the translation segments, a similar procedure was applied to the pool of 72 orientations. Each orientation in the pool was assigned an initial sampling weight of four (so the total initial weight across the pool was 72×4). On each sampling event the probability of selecting orientation i (P(i))was proportional to its current weight wi:

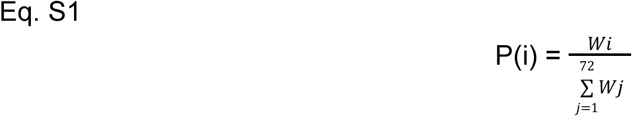

After an orientation was sampled its weight was reduced by 1, which lowered its probability of being chosen again. For each trial we selected three, four, or five orientations (for the target location of that trial), and the length of each translation segment was sampled uniformly from five to seven virtual meters. In addition, the above restrictions on the turn angle and bounded area applied. This way we ensured that the angular space is evenly sampled within the spatial constraints of the rectangular area.

**Figure S2.**
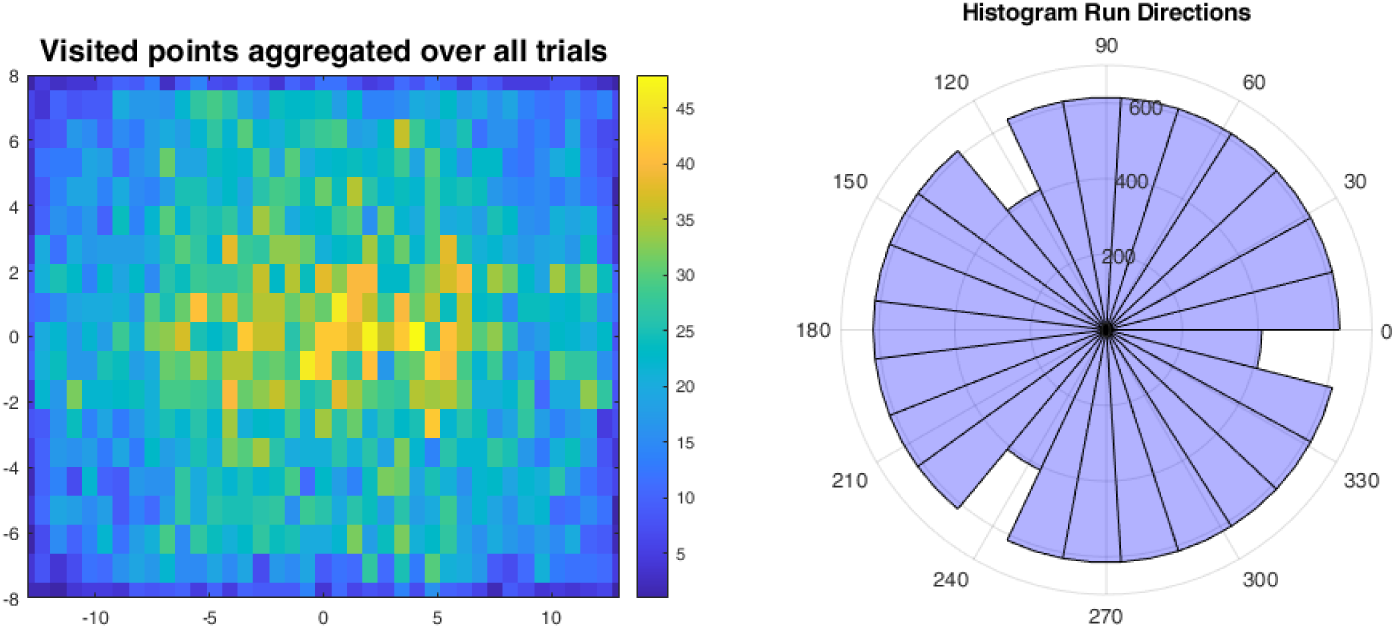
Left: spatial distribution of the waypoints presented across all trials and participants. Right: angular distribution of the orientation of translating trajectories across all trials and participants.

## Notes

### Competing Interest Statement

The authors have declared no competing interest.

## References

1. Barry, C., & Bush, D. (2012). From A to Z: A potential role for grid cells in spatial navigation. Neural Systems & Circuits, 2(1), 6. 10.1186/2042-1001-2-6

2. Bates, D., Mächler, M., Bolker, B., & Walker, S. (2015). Fitting Linear Mixed-Effects Models Using **lme4**. Journal of Statistical Software, 67(1). 10.18637/jss.v067.i01

3. Bellmund, J. L. S., de Cothi, W., Ruiter, T. A., Nau, M., Barry, C., & Doeller, C. F. (2020). Deforming the metric of cognitive maps distorts memory. Nature Human Behaviour, 4(2), 177–188. 10.1038/s41562-019-0767-3

4. Bellmund, J. L. S., Gärdenfors, P., Moser, E. I., & Doeller, C. F. (2018). Navigating cognition: Spatial codes for human thinking. Science, 362(6415), eaat6766. 10.1126/science.aat6766

5. Bigdely-Shamlo, N., Mullen, T., Kothe, C., Su, K.-M., & Robbins, K. A. (2015). The PREP pipeline: Standardized preprocessing for large-scale EEG analysis. Frontiers in Neuroinformatics, 9. 10.3389/fninf.2015.00016

6. Bush, D., Barry, C., Manson, D., & Burgess, N. (2015). Using Grid Cells for Navigation. Neuron, 87(3), 507–520. 10.1016/j.neuron.2015.07.006

7. Canolty, R. T., Edwards, E., Dalal, S. S., Soltani, M., Nagarajan, S. S., Kirsch, H. E., Berger, M. S., Barbaro, N. M., & Knight, R. T. (2006). High Gamma Power Is Phase-Locked to Theta Oscillations in Human Neocortex. Science, 313(5793), 1626–1628. 10.1126/science.1128115

8. Chen, D., Kunz, L., Lv, P., Zhang, H., Zhou, W., Liang, S., Axmacher, N., & Wang, L. (2021). Theta oscillations coordinate grid-like representations between ventromedial prefrontal and entorhinal cortex. Science Advances, 7(44), eabj0200. 10.1126/sciadv.abj0200

9. Chen, D., Kunz, L., Wang, W., Zhang, H., Wang, W.-X., Schulze-Bonhage, A., Reinacher, P. C., Zhou, W., Liang, S., Axmacher, N., & Wang, L. (2018). Hexadirectional Modulation of Theta Power in Human Entorhinal Cortex during Spatial Navigation. Current Biology, 28(20), 3310–3315.e4. 10.1016/j.cub.2018.08.029

10. Clark, H., & Nolan, M. F. (2024). Task-anchored grid cell firing is selectively associated with successful path integration-dependent behaviour. eLife, 12, RP89356. 10.7554/eLife.89356

11. Convertino, L., Bush, D., Zheng, F., Adams, R. A., & Burgess, N. (2023). Reduced grid-like theta modulation in schizophrenia. Brain, 146(5), 2191–2198. 10.1093/brain/awac416

12. de Cheveigné, A. (2020). ZapLine: A simple and effective method to remove power line artifacts. NeuroImage, 207, 116356. 10.1016/j.neuroimage.2019.116356

13. Debener, S., Minow, F., Emkes, R., Gandras, K., & de Vos, M. (2012). How about taking a low-cost, small, and wireless EEG for a walk? Psychophysiology, 49(11), 1617–1621. 10.1111/j.1469-8986.2012.01471.x

14. Delorme, A., & Makeig, S. (2004). EEGLAB: An open source toolbox for analysis of single-trial EEG dynamics including independent component analysis. Journal of Neuroscience Methods, 134(1), 9–21. 10.1016/j.jneumeth.2003.10.009

15. Doeller, C. F., Barry, C., & Burgess, N. (2010). Evidence for grid cells in a human memory network. Nature, 463(7281), 657–661. 10.1038/nature08704

16. Fiete, I. R., Burak, Y., & Brookings, T. (2008). What grid cells convey about rat location. Journal of Neuroscience, 28(27), 6858–6871. 10.1523/JNEUROSCI.5684-07.2008

17. Gil, M., Ancau, M., Schlesiger, M. I., Neitz, A., Allen, K., De Marco, R. J., & Monyer, H. (2018). Impaired path integration in mice with disrupted grid cell firing. Nature Neuroscience, 21(1), 81–91. 10.1038/s41593-017-0039-3

18. Ginosar, G., Aljadeff, J., Las, L., Derdikman, D., & Ulanovsky, N. (2023). Are grid cells used for navigation? On local metrics, subjective spaces, and black holes. Neuron, 111(12), 1858–1875. 10.1016/j.neuron.2023.03.027

19. Goeke, C., Kornpetpanee, S., Köster, M., Fernández-Revelles, A. B., Gramann, K., & König, P. (2015). Cultural background shapes spatial reference frame proclivity. Scientific Reports, 5(1), 11426. 10.1038/srep11426

20. Gorgolewski, K. J., Auer, T., Calhoun, V. D., Craddock, R. C., Das, S., Duff, E. P., Flandin, G., Ghosh, S. S., Glatard, T., Halchenko, Y. O., Handwerker, D. A., Hanke, M., Keator, D., Li, X., Michael, Z., Maumet, C., Nichols, B. N., Nichols, T. E., Pellman, J., … Poldrack, R. A. (2016). The brain imaging data structure, a format for organizing and describing outputs of neuroimaging experiments. Scientific Data, 3(1), 160044. 10.1038/sdata.2016.44

21. Gramann, K., Müller, H. J., Eick, E. M., & Schönebeck, B. (2005). Evidence of separable spatial representations in a virtual navigation task. Journal of Experimental Psychology: Human Perception and Performance, 31(6), 1199.

22. Gramann, K. (2013). Embodiment of Spatial Reference Frames and Individual Differences in Reference Frame Proclivity. Spatial Cognition & Computation, 13(1), 1–25. 10.1080/13875868.2011.589038

23. Gramann, K., Gwin, J. T., Ferris, D. P., Oie, K., Jung, T. P., Lin, C. T., Liao, L. D., & Makeig, S. (2011). Cognition in action: Imaging brain/body dynamics in mobile humans. https://opus.lib.uts.edu.au/handle/10453/116311

24. Gross, J., Kujala, J., Hämäläinen, M., Timmermann, L., Schnitzler, A., & Salmelin, R. (2001). Dynamic imaging of coherent sources: Studying neural interactions in the human brain. Proceedings of the National Academy of Sciences, 98(2), 694–699. 10.1073/pnas.98.2.694

25. Hafting, T., Fyhn, M., Molden, S., Moser, M.-B., & Moser, E. I. (2005). Microstructure of a spatial map in the entorhinal cortex. Nature, 436(7052), 801–806. 10.1038/nature03721

26. Hartley, T., Maguire, E. A., Spiers, H. J., & Burgess, N. (2003). The well-worn route and the path less traveled: distinct neural bases of route following and wayfinding in humans. Neuron, 37(5), 877–888.

27. He, Q., & Brown, T. I. (2019). Environmental Barriers Disrupt Grid-like Representations in Humans during Navigation. Current Biology, 29(16), 2718–2722.e3. 10.1016/j.cub.2019.06.072

28. Iggena, D., Jeung, S., Maier, P. M., Ploner, C. J., Gramann, K., & Finke, C. (2023). Multisensory input modulates memory-guided spatial navigation in humans. Communications Biology, 6(1), 1–15. 10.1038/s42003-023-05522-6

29. Jacobs, J., Miller, J., Lee, S. A., Coffey, T., Watrous, A. J., Sperling, M. R., Sharan, A., Worrell, G., Berry, B., Lega, B., Jobst, B. C., Davis, K., Gross, R. E., Sheth, S. A., Ezzyat, Y., Das, S. R., Stein, J., Gorniak, R., Kahana, M. J., & Rizzuto, D. S. (2016). Direct Electrical Stimulation of the Human Entorhinal Region and Hippocampus Impairs Memory. Neuron, 92(5), 983–990. 10.1016/j.neuron.2016.10.062

30. Jeung, S., Cockx, H., Appelhoff, S., Berg, T., Gramann, K., Grothkopp, S., Warmerdam, E., Hansen, C., Oostenveld, R., BIDS Maintainers, Appelhoff, S., Markiewicz, C. J., Salo, T., Gau, R., Blair, R., Galassi, A., Earl, E., Rogers, C., Hardcastle, N., … Welzel, J. (2024). Motion-BIDS: An extension to the brain imaging data structure to organize motion data for reproducible research. Scientific Data, 11(1), 716. 10.1038/s41597-024-03559-8

31. Julian, J. B., Keinath, A. T., Frazzetta, G., & Epstein, R. A. (2018). Human entorhinal cortex represents visual space using a boundary-anchored grid. Nature Neuroscience, 21(2), 191–194. 10.1038/s41593-017-0049-1

32. Jungnickel, E., Gehrke, L., Klug, M., & Gramann, K. (2019). Chapter 10—MoBI—Mobile Brain/Body Imaging. In H. Ayaz & F. Dehais (Eds), Neuroergonomics (pp. 59–63). Academic Press. 10.1016/B978-0-12-811926-6.00010-5

33. Klamer, S., Elshahabi, A., Lerche, H., Braun, C., Erb, M., Scheffler, K., & Focke, N. K. (2015). Differences Between MEG and High-Density EEG Source Localizations Using a Distributed Source Model in Comparison to fMRI. Brain Topography, 28(1), 87–94. 10.1007/s10548-014-0405-3

34. Klug, M., Jeung, S., Wunderlich, A., Gehrke, L., Protzak, J., Djebbara, Z., Argubi-Wollesen, A., Wollesen, B., & Gramann, K. (2022). The BeMoBIL Pipeline for automated analyses of multimodal mobile brain and body imaging data (p. 2022.09.29.510051). bioRxiv. 10.1101/2022.09.29.510051

35. Kothe, C., Shirazi, S. Y., Stenner, T., Medine, D., Boulay, C., Grivich, M. I., Mullen, T., Delorme, A., & Makeig, S. (2024). The Lab Streaming Layer for Synchronized Multimodal Recording. bioRxiv: The Preprint Server for Biology, 2024.02.13.580071. 10.1101/2024.02.13.580071

36. Krupic, J., Bauza, M., Burton, S., Barry, C., & O’Keefe, J. (2015). Grid cell symmetry is shaped by environmental geometry. Nature, 518(7538), 232–235. 10.1038/nature14153

37. Kunz, L., Schröder, T. N., Lee, H., Montag, C., Lachmann, B., Sariyska, R., Reuter, M., Stirnberg, R., Stöcker, T., Messing-Floeter, P. C., Fell, J., Doeller, C. F., & Axmacher, N. (2015). Reduced grid-cell-like representations in adults at genetic risk for Alzheimer’s disease. *Science (New York*, N.Y.), 350(6259), 430–433. 10.1126/science.aac8128

38. Lenth, R. V., Bolker, B., Buerkner, P., Giné-Vázquez, I., Herve, M., Jung, M., Love, J., Miguez, F., Piaskowski, J., Riebl, H., & Singmann, H. (2024). *emmeans: Estimated Marginal Means, aka Least-Squares Means* (Version 1.10.2) [Computer software]. https://cran.r-project.org/web/packages/emmeans/index.html

39. Maidenbaum, S., Miller, J., Stein, J. M., & Jacobs, J. (2018). Grid-like hexadirectional modulation of human entorhinal theta oscillations. Proceedings of the National Academy of Sciences, 115(42), 10798–10803. 10.1073/pnas.1805007115

40. Makeig, S., Gramann, K., Jung, T.-P., Sejnowski, T. J., & Poizner, H. (2009). Linking brain, mind and behavior. International Journal of Psychophysiology, 73(2), 95–100. 10.1016/j.ijpsycho.2008.11.008

41. Miller, J., Watrous, A. J., Tsitsiklis, M., Lee, S. A., Sheth, S. A., Schevon, C. A., Smith, E. H., Sperling, M. R., Sharan, A., Asadi-Pooya, A. A., Worrell, G. A., Meisenhelter, S., Inman, C. S., Davis, K. A., Lega, B., Wanda, P. A., Das, S. R., Stein, J. M., Gorniak, R., & Jacobs, J. (2018). Lateralized hippocampal oscillations underlie distinct aspects of human spatial memory and navigation. Nature Communications, 9(1), 2423. 10.1038/s41467-018-04847-9

42. Oostenveld, R., Fries, P., Maris, E., & Schoffelen, J.-M. (2011). FieldTrip: Open Source Software for Advanced Analysis of MEG, EEG, and Invasive Electrophysiological Data. Computational Intelligence and Neuroscience, 2011(1), 156869. 10.1155/2011/156869

43. Palmer, J. A., Kreutz-Delgado, K., & Makeig, S. (2011). AMICA: An Adaptive Mixture of Independent Component Analyzers with Shared Components.

44. Papale, A. E., Zielinski, M. C., Frank, L. M., Jadhav, S. P., & Redish, A. D. (2016). Interplay between Hippocampal Sharp-Wave-Ripple Events and Vicarious Trial and Error Behaviors in Decision Making. Neuron, 92(5), 975–982. 10.1016/j.neuron.2016.10.028

45. Park, S. A., Miller, D. S., & Boorman, E. D. (2021). Inferences on a multidimensional social hierarchy use a grid-like code. Nature Neuroscience, 24(9), 1292–1301. 10.1038/s41593-021-00916-3

46. Peng, J.-J., Throm, B., Najafian Jazi, M., Yen, T.-Y., Pizzarelli, R., Monyer, H., & Allen, K. (2025). Grid cells accurately track movement during path integration-based navigation despite switching reference frames. Nature Neuroscience, 28(10), 2092–2105. 10.1038/s41593-025-02054-6

47. Pernet, C. R., Appelhoff, S., Gorgolewski, K. J., Flandin, G., Phillips, C., Delorme, A., & Oostenveld, R. (2019). EEG-BIDS, an extension to the brain imaging data structure for electroencephalography. Scientific Data, 6(1), 103. 10.1038/s41597-019-0104-8

48. Pion-Tonachini, L., Kreutz-Delgado, K., & Makeig, S. (2019). ICLabel: An automated electroencephalographic independent component classifier, dataset, and website. NeuroImage, 198, 181–197. 10.1016/j.neuroimage.2019.05.026

49. Redish, A. D. (2016). Vicarious trial and error. Nature Reviews Neuroscience, 17(3), 147–159. 10.1038/nrn.2015.30

50. Robitaille, N., Marois, R., Todd, J., Grimault, S., Cheyne, D., & Jolicœur, P. (2010). Distinguishing between lateralized and nonlateralized brain activity associated with visual short-term memory: fMRI, MEG, and EEG evidence from the same observers. NeuroImage, 53(4), 1334–1345. 10.1016/j.neuroimage.2010.07.027

51. Rolls, E. T., Huang, C. C., Lin, C. P., Feng, J., & Joliot, M. (2020). Automated anatomical labelling atlas 3. Neuroimage, 206, 116189.

52. Schmidt, B., Papale, A., Redish, A. D., & Markus, E. J. (2013). Conflict between place and response navigation strategies: Effects on vicarious trial and error (VTE) behaviors. Learning & Memory, 20(3), 130–138. 10.1101/lm.028753.112

53. Stangl, M., Maoz, S. L., & Suthana, N. (2023). Mobile cognition: Imaging the human brain in the ‘real world’. Nature Reviews Neuroscience, 24(6), 347–362. 10.1038/s41583-023-00692-y

54. Staudigl, T., Leszczynski, M., Jacobs, J., Sheth, S. A., Schroeder, C. E., Jensen, O., & Doeller, C. F. (2018). Hexadirectional Modulation of High-Frequency Electrophysiological Activity in the Human Anterior Medial Temporal Lobe Maps Visual Space. Current Biology, 28(20), 3325–3329.e4. 10.1016/j.cub.2018.09.035

55. Styles, S. J., Ković, V., Ke, H., & Šoškić, A. (2021). Towards ARTEM-IS: Design guidelines for evidence-based EEG methodology reporting tools. NeuroImage, 245, 118721. 10.1016/j.neuroimage.2021.118721

56. Topalovic, U., Aghajan, Z. M., Villaroman, D., Hiller, S., Christov-Moore, L., Wishard, T. J., Stangl, M., Hasulak, N. R., Inman, C. S., Fields, T. A., Rao, V. R., Eliashiv, D., Fried, I., & Suthana, N. (2020). Wireless Programmable Recording and Stimulation of Deep Brain Activity in Freely Moving Humans. Neuron, 108(2), 322–334.e9. 10.1016/j.neuron.2020.08.021

57. Wang, N., Wang, Y., Guo, M., Wang, L., Wang, X., Zhu, N., Yang, J., Wang, L., Zheng, C., & Ming, D. (2025). Dynamic gamma modulation of hippocampal place cells predominates development of theta sequences. eLife, 13, RP97334. 10.7554/eLife.97334

58. Wang, W., & Wang, W. (2021). Effect of reward on electrophysiological signatures of grid cell population activity in human spatial navigation. Scientific Reports, 11(1), 23577. 10.1038/s41598-021-03124-y

59. White, D. J., Congedo, M., Ciorciari, J., & Silberstein, R. B. (2012). Brain oscillatory activity during spatial navigation: Theta and gamma activity link medial temporal and parietal regions. Journal of Cognitive Neuroscience, 24(3), 686–697. 10.1162/jocn_a_00098

